# Relation of in-utero exposure to antiepileptic drugs to pregnancy duration and size at birth

**DOI:** 10.1101/574269

**Authors:** Andrea V Margulis, Sonia Hernandez-Diaz, Thomas McElrath, Kenneth J Rothman, Estel Plana, Catarina Almqvist, Brian M D’Onofrio, Anna Sara Oberg

## Abstract

**Background:** The associations of individual antiepileptic drugs (AEDs) with pregnancy duration and size at birth, and potential dose relations, are not well characterized.

**Methods:** This cohort study used nationwide Swedish register data (1996-2013). Adjusting for smoking, epilepsy and other AED indications, we used linear and quantile regression to explore associations with pregnancy duration, and birth weight, length, and head circumference (the last three operationalized as z-scores). We used logistic regression for preterm delivery, small for gestational age, and microcephaly. Lamotrigine was the reference drug.

**Results:** 6,720 infants were exposed to AEDs in utero; AED exposure increased over the study period. Relative to lamotrigine-exposed infants, carbamazepine-exposed infants were born, on average, 1.3 days earlier (mean [95% confidence interval]: −1.3 [−2.3 to −0.3]); were 0.1 standard deviations (SDs) lighter (−0.1 [−0.2 to 0.0]); and had a head circumference that was 0.2 SDs smaller (−0.2 [−0.3 to −0.1]). Pregabalin-exposed infants were born, on average, 1.1 days earlier (-1.1 [−3.0 to 0.8]); were 0.1 SDs lighter (−0.1 [−0.3 to 0.0]); and had the same head circumference. Levetiracetam-exposed infants were born, on average, 0.5 days earlier (−0.5 [−2.6 to 1.6]); were 0. 1 SDs lighter (−0.1 [−0.3 to 0.0]); and were 0.1 SDs smaller (−0.1 [−0.3 to 0.1]) in head circumference. Valproic acid–exposed infants had, on average, the same duration of gestation and birth weight z-score, but were 0.2 SDs smaller (−0.2 [−0.2 to −0.1]) in head circumference. More negative associations at the left tail of pregnancy duration and birth weight z-score, effect-measure modification, and dose-response relations were noted for some of the associations. Observed associations were generally of smaller magnitude than that of smoking, assessed as a potential confounder in the same models.

**Conclusions:** In comparison with lamotrigine, valproic acid and carbamazepine had a more negative association with head circumference than other study AEDs.

## INTRODUCTION

Epilepsy and antiepileptic drugs (AEDs) have been associated with adverse pregnancy, fetal, and neonatal outcomes [1]. AEDs differ in their risk for congenital malformations [2–4], and some associations have been found to be dose dependent [4–6]. Newer AEDs are generally considered safer than the older drugs, with the possible exception of topiramate [7]. Antiepileptic drugs also differ in the magnitude of their associations with adverse neurodevelopmental outcomes in the offspring, which also appear to be dose dependent [8–10]. The exploration of indication and dose is important because confounding by indication has been a concern and AED doses are often higher in epilepsy than in other conditions [11].

A meta-analysis has shown elevated point estimates for the association of AEDs, as a group, with shortened pregnancies and reduced birth size [1], but comparative safety evidence for these endpoints is scarce, as demonstrated by a systematic literature search we conducted to inform our decision on which AED to use as a reference drug [12] and to provide context to the present study. We identified 15 papers that provided adjusted comparisons for individual AEDs [13–27], of which 12 used unexposed populations as the reference (details on this literature search are in Supporting Information file 1).

Furthermore, previous research has assessed associations with binarized endpoints or associations only at the mean of the continuous distributions. In this study, we sought to explore the comparative safety of individual AEDs on pregnancy duration and birth weight, length, and head circumference and to explore dose relations on these endpoints, adjusting for epilepsy and other indications. To characterize effects thoroughly, we assessed continuous and binary forms of the endpoints and investigated potential AED effects in both tails of the endpoint distributions. Advantages of this comparative safety design, in which we used lamotrigine as the reference instead of no AED use, are that confounding by indication is partially removed and that study results will better inform the choice of patients and clinicians when antiepileptic treatment is needed.

## METHODS

### Overview

We conducted a cohort study based on nationwide Swedish register data from 1996 through 2013 to explore the association between maternal use of individual AEDs and pregnancy duration and fetal size. Lamotrigine was the reference AED because it is commonly used and has been considered to have fewer adverse fetal effects than other AEDs [2, 12, 28, 29].

### Data sources

In Sweden, tax-funded health care is provided to all citizens. Information arising from contacts with the health care system is collected in registries that can be linked through a unique personal registration number assigned to all individuals residing in Sweden. Drugs are coded in the Anatomic Therapeutic Chemical classification system, and diagnoses are coded using the International Classification of Diseases (10th revision since 1997).

The Swedish Medical Birth Register [30] collects information from prenatal care, including selfreported medication use at first and subsequent visits, and from standardized delivery charts, including gestational age at birth, birth weight, length, and head circumference. Information on medication use in the first visit is more complete than in subsequent visits. Medications noted only in free-text comments have been coded and incorporated in the structured drug fields. The Prescribed Drug Register records all prescription medications dispensed by pharmacies since 1 July 2005. Information available from prescriptions include drug name, drug strength, number of packages dispensed, and number of defined daily doses (DDDs) per package [31]. The National Patient Register includes all discharge records from hospitalizations since 1987 and 75%-80% of visits to specialists, including psychiatric care, since 2001. The Swedish Register of Education contains information on the maximum education level attained per year [32]. The Total Population Register contains demographic and administrative information including nationality and birth and migration dates [33].

### Study population

The study population included all women with records for AEDs in pregnancy who delivered a live infant with gestational age of 24 to 42 completed weeks in 1996-2013 and their newborns. Infants born from women who immigrated less than 12 months before pregnancy and infants with chromosomal abnormalities were excluded. Infants with congenital malformations and no chromosomal abnormalities and infants from multiple pregnancies were included. All eligible infants per woman were included.

### Exposure

We report on the five AEDs that were most commonly used in pregnancy in the last year of our study period: carbamazepine, valproic acid, pregabalin, levetiracetam, and lamotrigine. We defined three exposure windows for analysis: any time in pregnancy, first trimester (regardless of whether treatment was later discontinued), and first and second/third trimesters (“continuers”).

To create the exposure variables, information on first-trimester exposure was obtained from prescriptions dispensed between the first day of the last menstrual period and gestational day 89 and from self-report in the first prenatal visit in women who started prenatal care by gestational week 15. Information on second-/third-trimester exposure was obtained from prescriptions dispensed between day 90 and the day before delivery, from self-reports in the first prenatal visit in women who started prenatal care after gestational week 15, and from self-reports in subsequent prenatal visits (self-reports did not allow a clear differentiation of second-versus third-trimester exposure; thus, we combined both periods). Because of incomplete capture of self-reports after the first prenatal visit, exposure in continuers was defined only for the period for which dispensing data were available (deliveries in 2006-2013). Women and infants exposed to more than one AED were considered to be exposed to each of them.

Dose was derived from dispensed prescriptions (deliveries in 2006-2013). For each prescription, dose was calculated by multiplying the number of packs dispensed by the number of DDDs per pack and by the number of milligrams in a DDD [31]. The mean daily dose was calculated separately for each AED per infant by dividing the dose in prescriptions dispensed between the first day of the last menstrual period and the day before delivery over the number of days in the same period.

### Characteristics of the study population

We extracted medical and obstetric information from the national health registers, which derive their information from prenatal care records, hospitalization records, outpatient specialist care records, and dispensed prescriptions. Codes, source of data, timing of ascertainment, categorization, and other details for medical and other characteristics are presented in Supporting information file 2.

### Endpoints

Study endpoints were duration of pregnancy, preterm delivery, birth weight, small for gestational age (SGA), length at birth, head circumference at birth, and microcephaly, all ascertained from the Medical Birth Register. Duration of pregnancy is predominantly based on ultrasound estimation [34] and is recorded in days; preterm delivery was defined as delivery before 37 completed weeks. Birth weight, length, and head circumference were operationalized as z-scores to assess size independently from gestational age at birth; the birth weight z-score for each infant is the observed birth weight minus the reference mean birth weight, divided by the reference birth weight standard deviation (SD), where the mean and SD were those for infants born at the same gestational age, using a local standard [35]. Small for gestational age was defined within the Medical Birth Register from standard growth curves based on ultrasound-derived fetal weights for singletons only [36]. Microcephaly was defined within the Medical Birth Register as a head circumference of two or more SDs below the mean for gestational age at birth, using a local standard [35].

### Statistical analyses

In the main analysis, continuous endpoints were analyzed using linear regression and quantile regression for the 10th, 50th, and 90th percentiles [37]. Lamotrigine was the reference drug. We produced unadjusted results and results adjusted for maternal age at delivery, education, country of origin, marital status, early pregnancy body mass index, smoking in current pregnancy, alcohol dependence, diabetes, hypertension, epilepsy, depression, bipolar disorder, migraine, chronic pain, other psychiatric disorders, and year of delivery. Variable definitions are presented in Supporting information file 2. Missing values (Table 1) were imputed for analysis as the most commonly observed value in the study population; multiple imputation had been planned for variables with missing values in 10% or more of the observations, but missingness was below that threshold. Binary endpoints were analyzed using logistic regression. We conducted adjusted analyses in comparisons with five or more events in the smallest cell (i.e., exposed cases, exposed noncases, unexposed cases, unexposed noncases), adjusting for the variables listed above. We used the weighted copy method to facilitate the convergence of logistic regression models. With this method, analyses are conducted on an expanded data set that consists of the original data set and a copy of the data with the outcomes reversed; confidence intervals are adjusted by the use of weights in the code [38–40]. We weighted the original data 999 times that of the reversed data. The unit of analysis was pregnancy for the endpoints duration of pregnancy and preterm delivery; for other endpoints, the unit of analysis was infant.

**Table 1.** Characteristics of Study Population and Mean Daily Dose By Antiepileptic Drug

| Characteristic | Lamotrigine | Carbamazepine | Pregabalin | Levetiracetam | Valproic acid |
| --- | --- | --- | --- | --- | --- |
| Number of exposed women | 1,757 | 1,529 | 542 | 245 | 809 |
| Number of exposed infants | 2,254 | 2,095 | 562 | 307 | 1,137 |
| Age at delivery (years) |  |  |  |  |  |
| 24 or less | 427 (18.9%) | 249 (11.9%) | 119 (21.2%) | 52 (16.9%) | 206 (18.1%) |
| 25-29 | 691 (30.7%) | 626 (29.9%) | 160 (28.5%) | 92 (30.0%) | 350 (30.8%) |
| 30-34 | 721 (32.0%) | 728 (34.7%) | 151 (26.9%) | 114 (37.1%) | 374 (32.9%) |
| 35 or more | 415 (18.4%) | 492 (23.5%) | 132 (23.5%) | 49 (16.0%) | 207 (18.2%) |
| Mother's country of origin |  |  |  |  |  |
| Nordic countries | 2,040 (90.5%) | 1,812 (86.5%) | 486 (86.5%) | 257 (83.7%) | 995 (87.5%) |
| Other European countries | 86 (3.8%) | 72 (3.4%) | 21 (3.7%) | 16 (5.2%) | 59 (5.2%) |
| Asia | 82 (3.6%) | 128 (6.1%) | 39 (6.9%) | 24 (7.8%) | 60 (5.3%) |
| Others | 46 (2.0%) | 83 (4.0%) | 16 (2.8%) | 10 (3.3%) | 23 (2.0%) |
| Maternal education |  |  |  |  |  |
| Up to 12 years | 1,396 (61.9%) | 1,340 (64.0%) | 461 (82.0%) | 182 (59.3%) | 759 (66.8%) |
| 13 years or more | 826 (36.6%) | 717 (34.2%) | 96 (17.1%) | 115 (37.5%) | 364 (32.0%) |
| No information | 32 (1.4%) | 38 (1.8%) | 5 (0.9%) | 10 (3.3%) | 14 (1.2%) |
| Maternal marital status |  |  |  |  |  |
| Lives with child's father | 1,926 (85.4%) | 1,854 (88.5%) | 382 (68.0%) | 275 (89.6%) | 1,004 (88.3%) |
| Does not live with child's father | 253 (11.2%) | 184 (8.8%) | 157 (27.9%) | 25 (8.1%) | 93 (8.2%) |
| No information | 75 (3.3%) | 57 (2.7%) | 23 (4.1%) | 7 (2.3%) | 40 (3.5%) |
| Early pregnancy BMI (kg/m<sup>2</sup>) |  |  |  |  |  |
| Less than 18.5 | 44 (2.0%) | 33 (1.6%) | 9 (1.6%) | 8 (2.6%) | 22 (1.9%) |
| 18.5 to less than 25 | 1,097 (48.7%) | 1,066 (50.9%) | 243 (43.2%) | 174 (56.7%) | 513 (45.1%) |
| 25 to less than 30 | 595 (26.4%) | 500 (23.9%) | 145 (25.8%) | 72 (23.5%) | 318 (28.0%) |
| 30 or more | 341 (15.1%) | 287 (13.7%) | 120 (21.4%) | 36 (11.7%) | 179 (15.7%) |
| Obese (codes for obesity) | 2 (0.1%) | 2 (0.1%) | 2 (0.4%) | 0 (0.0%) | 2 (0.2%) |
| No information | 175 (7.8%) | 207 (9.9%) | 43 (7.7%) | 17 (5.5%) | 103 (9.1%) |
| Smoking during pregnancy |  |  |  |  |  |
| Smoker | 386 (17.1%) | 258 (12.3%) | 234 (41.6%) | 26 (8.5%) | 211 (18.6%) |
| Nonsmoker | 1,809 (80.3%) | 1,787 (85.3%) | 310 (55.2%) | 277 (90.2%) | 894 (78.6%) |
| No information | 59 (2.6%) | 50 (2.4%) | 18 (3.2%) | 4 (1.3%) | 32 (2.8%) |
| Alcohol dependence | 123 (5.5%) | 38 (1.8%) | 69 (12.3%) | 1 (0.3%) | 25 (2.2%) |
| AED indications/uses |  |  |  |  |  |
| Epilepsy | 1,559 (69.2%) | 1,774 (84.7%) | 37 (6.6%) | 303 (98.7%) | 939 (82.6%) |
| Depression | 445 (19.7%) | 106 (5.1%) | 273 (48.6%) | 19 (6.2%) | 86 (7.6%) |
| Bipolar disorder | 460 (20.4%) | 27 (1.3%) | 57 (10.1%) | 2 (0.7%) | 74 (6.5%) |
| Other psychiatric disorders | 645 (28.6%) | 179 (8.5%) | 371 (66.0%) | 37 (12.1%) | 162 (14.2%) |
| Migraine | 224 (9.9%) | 105 (5.0%) | 132 (23.5%) | 24 (7.8%) | 62 (5.5%) |
| Chronic pain | 575 (25.5%) | 283 (13.5%) | 387 (68.9%) | 78 (25.4%) | 172 (15.1%) |
| Restless legs syndrome | 9 (0.4%) | 6 (0.3%) | 13 (2.3%) | 1 (0.3%) | 2 (0.2%) |
| None of the above | 63 (2.8%) | 222 (10.6%) | 30 (5.3%) | 3 (1.0%) | 93 (8.2%) |
| Diabetes | 76 (3.4%) | 56 (2.7%) | 24 (4.3%) | 10 (3.3%) | 43 (3.8%) |
| Hypertension | 100 (4.4%) | 100 (4.8%) | 48 (8.5%) | 12 (3.9%) | 65 (5.7%) |
| Medications in current pregnancy |  |  |  |  |  |
| AED polytherapy | 442 (19.6%) | 269 (12.8%) | 63 (11.2%) | 180 (58.6%) | 256 (22.5%) |
| Antidepressants | 458 (20.3%) | 107 (5.1%) | 284 (50.5%) | 15 (4.9%) | 117 (10.3%) |
| Antipsychotics | 157 (7.0%) | 31 (1.5%) | 76 (13.5%) | 3 (1.0%) | 50 (4.4%) |
| Migraine treatment | 51 (2.3%) | 21 (1.0%) | 43 (7.7%) | 6 (2.0%) | 21 (1.8%) |
| Opioids | 165 (7.3%) | 79 (3.8%) | 195 (34.7%) | 27 (8.8%) | 55 (4.8%) |
| Female infant | 1,178 (52.3%) | 982 (46.9%) | 285 (50.7%) | 144 (46.9%) | 557 (49.0%) |
| High dose (mean, mg/day)<sup>a</sup> | 454 | 905 | 384 | 2,489 | 1,349 |
| Low dose (mean, mg/day)<sup>b</sup> | 41 | 186 | 12 | 402 | 211 |

AED, antiepileptic drug; BMI, body mass index.
Note = The denominator for calculations is the number of infants.
<sup>a</sup> Mean dose in top tertile of pregnancy daily dose.
<sup>b</sup> Mean dose in bottom tertile of pregnancy daily dose.

The Regional Ethical Review Board in Stockholm, Sweden, approved the linkage of registers to perform this type of study (DNR 2013/862-31/5). This study was judged to be exempt from review by the RTI International institutional review board.

As secondary and sensitivity analyses, to better understand the influence of the underlying maternal health problem being treated, we repeated the main analysis in mothers with a diagnosis of epilepsy or chronic pain. We also explored the influence of monotherapy versus polytherapy (e.g., carbamazepine in polytherapy [not including lamotrigine] vs. lamotrigine in polytherapy [not including carbamazepine]). To address potential exposure misclassification and biases related to missing data, we conducted analyses on women with definite exposure (women in whom AED use from self-reports and dispensed prescriptions were consistent) and a complete case analysis. Addressing whether associations might be driven by in-utero crowding or malformations, we repeated analyses in singletons with no major congenital malformations. We repeated the analyses in the first pregnancy or infant per woman to gain understanding on any statistical effect of ignoring the correlation among siblings. We also explored associations separately in female and male infants. We explored effect-measure modification separately by smoking and use of selective serotonin reuptake inhibitors (SSRIs) in pregnancy in linear regression analyses by incorporating an appropriate interaction term into the regression models.

In dose analyses, we compared the top tertile of mean daily dose with the bottom tertile (which served as the reference) for each individual AED, using linear regression. All models were adjusted as in the main analysis, and the weighted copy method was used for binary endpoints.

We present results from a subset of analyses in the body of this paper; others, including analyses on birth weight, length, and head circumference as recorded (in grams or centimeters, as opposed to z-scores), are included in Supporting information file 3 (Tables S1-S9).

## RESULTS

### Study population

The study population comprised 6,720 infants born to 5,112 women. Antiepileptic drug use in pregnancy increased from 181 exposed infants in 1996 to 607 in 2013 (Figure 1). In 2013, the most commonly used AEDs were lamotrigine (47%), carbamazepine (16%), pregabalin (16%), levetiracetam (10%), and valproic acid (8%); we present results on these drugs. The prevalences of most maternal characteristics were quite homogeneous across users of individual study AEDs (Table 1), except for the medical conditions for which study AEDs are prescribed.

**Figure 1.**
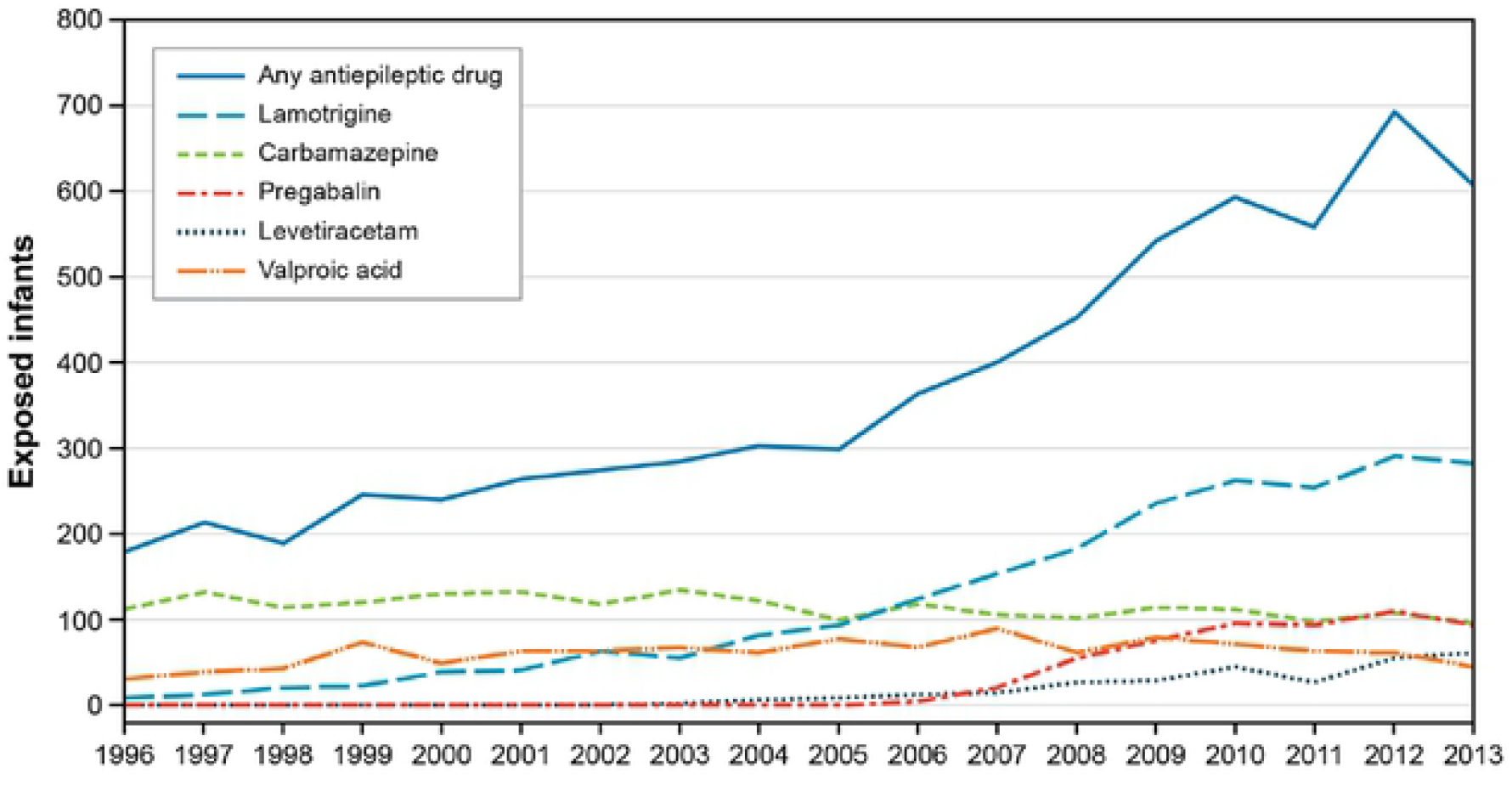
Use of antiepileptic drugs in pregnancy, Sweden 1996-2013. Note = Year represents year of delivery. The curve labeled “any antiepileptic drug” includes all drugs in chapter N03 of the Anatomical Therapeutic Chemical (ATC) classification system.

### Carbamazepine

Carbamazepine use decreased over the study period from 63% of AED-exposed infants in 1996 to 16% in 2013 (Figure 1); mothers of 85% of carbamazepine-exposed infants had an epilepsy diagnosis, and 13% of infants were exposed to AED polytherapy (Table 1).

We observed a pattern of slightly shorter pregnancies with linear regression models (mean [95% confidence interval]: −1.3 [−2.3 to −0.3] days) and smaller infants after exposure to carbamazepine, relative to lamotrigine, with an asymmetrical effect in which the head circumference z-score was somewhat more affected (−0.2 [−0.3 to −0.1] SDs) than birth weight or birth length z-scores (both at −0.1 [−0.2 to 0] SDs) (Table 2 and Supporting information file 3, Table S1). Associations at the 10th percentile of pregnancy duration were generally more negative than associations at the 90th percentile (i.e., regression coefficients from quantile regression models for carbamazepine indicated that exposure to carbamazepine was associated with a shorter pregnancy duration when assessed at the 10th percentile of pregnancy duration than when assessed at the 90th percentile). Most odds ratios (ORs) from logistic regression models for preterm delivery, SGA, and microcephaly ranged between 1.1 and 1.5; observed effects were larger in infants exposed to polytherapy. Odds ratios for SGA and microcephaly in women with chronic pain were also larger. Exposure to SSRIs operated as an effect-measure modifier for duration of gestation, with shorter pregnancies (mean −5.8 [−9.7 to −2.0] days) in women exposed to both carbamazepine and SSRIs (Supporting information file 3, Table S2). High doses of carbamazepine were associated with higher risk for all outcomes relative to low doses of carbamazepine (Table 2 and Supporting information file 3, Table S1).

**Table 2.** Association Between in-Utero Carbamazepine Exposure and the Endpoints Duration of Pregnancy and Size at Birth

|  | Exposed to<br>Carbamazepin<br>e/Reference,<br>n/n | Difference (95% CI) |  |  |  | Odds Ratio<br>(95% CI) |
| --- | --- | --- | --- | --- | --- | --- |
|  |  | Percentile |  |  |  |  |
|  |  | Mean | 10th | 50th | 90th |  |
|  |  | Pregnancy duration (days) |  |  |  | Preterm birth |
| Use any time in pregnancy, carbamazepine vs. lamotrigine | 1,975 / 2,123 | -1.3 (-2.3 to -0.3) | -1.1 (-3.1 to 0.9) | -0.9 (-1.8 to 0.1) | -0.1 (-1.3 to 1.0) | 1.2 (0.9 to 1.5) |
| Use in first trimester, carbamazepine vs. lamotrigine | 1,686 / 1,930 | -1.6 (-2.7 to -0.5) | -2.3 (-4.5 to -0.1) | -0.9 (-1.8 to 0.0) | -0.5 (-1.5 to 0.6) | 1.3 (1.0 to 1.8) |
| Continuers, carbamazepine vs. lamotrigine | 459 / 1,013 | -1.3 (-3.0 to 0.3) | 0.0 (-3.8 to 3.8) | -0.3 (-2.0 to 1.3) | -0.5 (-1.9 to 0.9) | 1.1 (0.7 to 1.7) |
| Mother with epilepsy, carbamazepine vs. lamotrigine | 1,665 / 1,447 | -1.3 (-2.4 to -0.2) | -1.6 (-3.5 to 0.3) | -0.5 (-1.5 to 0.5) | -0.2 (-1.3 to 0.9) | 1.3 (0.9 to 1.7) |
| Mother with chronic pain, carbamazepine vs. lamotrigine | 259 / 541 | -1.5 (-4.2 to 1.1) | -4.5 (-10.5 to 1.5) | -0.7 (-2.7 to 1.4) | 0.1 (-2.7 to 2.8) | 1.3 (0.7 to 2.3) |
| Polytherapy, carbamazepine vs. lamotrigine | 167 / 336 | -2.4 (-5.8 to 1.0) | -6.1 (-15.1 to 2.8) | -2.0 (-5.4 to 1.4) | -1.5 (-4.0 to 1.0) | 1.7 (0.9 to 3.3) |
| High vs. low dose of carbamazepine | 264 / 275 | -4.6 (-7.5 to -1.6) | -6.8 (-12.6 to -0.9) | -3.4 (-5.8 to -0.9) | -2.1 (-4.7 to 0.4) | 2.8 (1.3 to 6.0) |
|  |  | Birth weight z-score |  |  |  | SGA |
| Use any time in pregnancy, carbamazepine vs. lamotrigine | 1,988 / 2,147 | -0.1 (-0.2 to -0.0) | -0.0 (-0.1 to 0.1) | -0.1 (-0.2 to -0.0) | -0.2 (-0.3 to -0.1) | 1.4 (0.9 to 2.1) |
| Use in first trimester, carbamazepine vs. lamotrigine | 1,699 / 1,953 | -0.1 (-0.2 to -0.0) | -0.1 (-0.2 to 0.1) | -0.1 (-0.2 to -0.0) | -0.2 (-0.3 to -0.1) | 1.7 (1.0 to 2.6) |
| Continuers, carbamazepine vs. lamotrigine | 466 / 1,021 | -0.1 (-0.2 to -0.0) | -0.1 (-0.3 to 0.1) | -0.1 (-0.2 to 0.0) | -0.2 (-0.3 to -0.0) | 1.3 (0.7 to 2.6) |
| Mother with epilepsy, carbamazepine vs. lamotrigine | 1,676 / 1,459 | -0.1 (-0.2 to -0.0) | 0.0 (-0.1 to 0.1) | -0.1 (-0.2 to -0.1) | -0.2 (-0.3 to -0.0) | 1.2 (0.8 to 1.9) |
| Mother with chronic pain, carbamazepine vs. lamotrigine | 263 / 552 | -0.2 (-0.3 to 0.0) | -0.1 (-0.4 to 0.3) | -0.1 (-0.3 to 0.1) | -0.1 (-0.4 to 0.1) | 1.8 (0.8 to 4.2) |
|  |  | Percentile |  |  |  |  |
|  |  | Mean | 10th | 50th | 90th |  |
| Polytherapy, carbamazepine vs. lamotrigine | 167 / 339 | -0.5 (-0.7 to -0.3) | -0.6 (-0.9 to -0.3) | -0.5 (-0.7 to -0.2) | -0.3 (-0.8 to 0.1) | 4.2 (1.2 to 14.4) |
| High vs. low dose of carbamazepine | 267 / 275 | -0.1 (-0.3 to 0.1) | -0.1 (-0.4 to 0.2) | -0.1 (-0.3 to 0.1) | -0.1 (-0.4 to 0.1) | 2.0 (0.7 to 5.6) |
| Birth length z-score |  |  |  |  |  |  |
| Use any time in pregnancy, carbamazepine vs. lamotrigine | 1,963 / 2,119 | -0.1 (-0.2 to -0.0) | -0.1 (-0.2 to 0.0) | -0.1 (-0.2 to 0.0) | -0.2 (-0.3 to -0.0) |  |
| Use in first trimester, carbamazepine vs. lamotrigine | 1,681 / 1,930 | -0.1 (-0.2 to -0.0) | -0.1 (-0.2 to 0.0) | -0.1 (-0.2 to -0.0) | -0.2 (-0.3 to -0.1) |  |
| Continuers, carbamazepine vs. lamotrigine | 461 / 1,006 | -0.2 (-0.3 to -0.1) | -0.2 (-0.4 to 0.0) | -0.2 (-0.3 to -0.1) | -0.2 (-0.4 to -0.1) |  |
| Mother with epilepsy, carbamazepine vs. lamotrigine | 1,655 / 1,441 | -0.1 (-0.2 to -0.0) | -0.1 (-0.3 to -0.0) | -0.1 (-0.2 to 0.0) | -0.1 (-0.3 to -0.0) |  |
| Mother with chronic pain, carbamazepine vs. lamotrigine | 260 / 542 | -0.2 (-0.4 to -0.0) | -0.1 (-0.3 to 0.2) | -0.2 (-0.4 to 0.1) | -0.3 (-0.6 to -0.0) |  |
| Polytherapy, carbamazepine vs. lamotrigine | 163 / 331 | -0.3 (-0.5 to -0.1) | -0.2 (-0.5 to 0.1) | -0.2 (-0.5 to 0.0) | -0.6 (-0.8 to -0.3) |  |
| High vs. low dose of carbamazepine | 260 / 273 | -0.1 (-0.3 to 0.0) | -0.1 (-0.4 to 0.1) | -0.2 (-0.4 to -0.0) | -0.0 (-0.3 to 0.2) |  |
| Birth head circumference z-score |  |  |  |  |  | Microcephaly |
| Use any time in pregnancy, carbamazepine vs. lamotrigine | 1,883 / 2,096 | -0.2 (-0.3 to -0.1) | -0.2 (-0.3 to -0.0) | -0.2 (-0.3 to -0.1) | -0.2 (-0.3 to -0.1) | 1.2 (0.7 to 1.9) |
| Use in first trimester, carbamazepine vs. lamotrigine | 1,605 / 1,906 | -0.2 (-0.3 to -0.2) | -0.2 (-0.4 to -0.1) | -0.3 (-0.4 to -0.2) | -0.3 (-0.4 to -0.2) | 1.3 (0.8 to 2.1) |
| Continuers, carbamazepine vs. lamotrigine | 456 / 1,002 | -0.3 (-0.4 to -0.2) | -0.3 (-0.5 to -0.1) | -0.4 (-0.5 to -0.2) | -0.4 (-0.6 to -0.2) | 1.3 (0.6 to 3.3) |
| Mother with epilepsy, carbamazepine vs. lamotrigine | 1,585 / 1,421 | -0.2 (-0.3 to -0.1) | -0.2 (-0.4 to -0.1) | -0.2 (-0.3 to -0.1) | -0.2 (-0.3 to -0.1) | 1.2 (0.7 to 1.9) |
| Mother with chronic pain, carbamazepine vs. lamotrigine | 256 / 543 | -0.2 (-0.4 to -0.0) | -0.2 (-0.6 to 0.1) | -0.2 (-0.4 to -0.1) | 0.0 (-0.2 to 0.3) | 2.7 (0.8 to 9.1) |
| Polytherapy, carbamazepine vs. lamotrigine | 155 / 329 | -0.6 (-0.8 to -0.4) | -0.5 (-0.8 to -0.2) | -0.6 (-0.8 to -0.3) | -0.7 (-1.0 to -0.4) | 2.6 (0.9 to 7.3) |
|  |  | Mean | Percentile |  |  |  |
|  |  |  | 10th | 50th | 90th |  |
| High vs. low dose of carbamazepine | 256 / 271 | -0.2 (-0.4 to 0.0) | -0.2 (-0.6 to 0.1) | -0.3 (-0.6 to -0.0) | -0.2 (-0.5 to 0.1) | Not applicable |

AED, antiepileptic drug; CI, confidence interval; SGA, small for gestational age.
Note = AED use was ascertained at any time in pregnancy, except where noted (indented rows). Analyses on continuers used data from deliveries in 2006-2013. In analyses of carbamazepine vs. lamotrigine, the reference was lamotrigine in the same exposure window. In dose-response analyses, the reference was the bottom tertile of mean daily dose of carbamazepine (2006-2013). All results were adjusted for birth year, maternal age at delivery, education, country of origin, marital status, body mass index, smoking in current pregnancy, alcohol dependence, diabetes, hypertension, epilepsy, depression, bipolar disorder, migraine, chronic pain, and other psychiatric disorders. When the smallest cell count was < 5, we did not produce adjusted results ("not applicable"). Models restricted to polytherapy compared infants exposed to carbamazepine and another AED (except lamotrigine) with those exposed to lamotrigine and another AED (except carbamazepine).

### Pregabalin

Despite first appearing in 2006, pregabalin was the third most commonly used AED in this cohort in 2013 (16% of infants). Pregabalin users differed from users of other AEDs: pregabalin users were younger (and had fewer years of education), lived less frequently with the infant’s father, and were more likely to be obese or smokers. Chronic pain was common among mothers of pregabalin-exposed infants (69% of pregabalin-exposed infants), as were psychiatric conditions comprising psychoses, panic attacks, and other conditions (“other psychiatric disorders” in Table 1, 66%); mothers of 7% of infants had an epilepsy diagnosis, and mothers of 11% were on AED polytherapy (Table 1).

Pregabalin-exposed pregnancies were slightly shorter than lamotrigine-exposed pregnancies (−1.1 [−3.0 to 0.8] days on average), which was more notable in women with a diagnosis of epilepsy (−5.6 [−10.7 to −0.4] days on average) (Table 3 and Supporting information file 3, Table S3). Birth weight and length z-scores were slightly smaller in pregabalin-exposed than in lamotrigine-exposed infants (−0.1 [−0.3 to 0] and −0.1 [−0.2 to 0] SDs on average, respectively), and head circumference z-score was less affected (0 [−0.1 to 0.1] SDs on average). Among continuers, though, the OR for microcephaly was 5.3 (0.9 to 30.8). The association with pregnancy duration appeared to be more pronounced when the fetus was female, while the opposite was true for head circumference. No clear effect-measure modification with smoking or SSRI use, and no dose-response relation were observed (Supporting information file 3, Table S4).

**Table 3.** Association Between in-Utero Pregabalin Exposure and the Endpoints Duration of Pregnancy and Size at Birth

|  | Exposed to<br>Pregabalin/Ref<br>erence, n/n | Difference (95% CI) |  |  |  | Odds Ratio<br>(95% CI) |
| --- | --- | --- | --- | --- | --- | --- |
|  |  | Percentile |  |  |  |  |
|  |  | Mean | 10th | 50th | 90th |  |
|  |  | Pregnancy duration (days) |  |  |  | Preterm birth |
| Use any time in pregnancy, pregabalin vs. lamotrigine | 522 / 2,190 | -1.1 (-3.0 to 0.8) | -2.7 (-6.7 to 1.2) | -0.5 (-2.5 to 1.4) | 0.3 (-1.6 to 2.3) | 1.5 (1.0 to 2.4) |
| Use in first trimester, pregabalin vs. lamotrigine | 484 / 1,977 | -1.8 (-3.7 to 0.2) | -3.2 (-7.5 to 1.1) | -0.5 (-2.4 to 1.3) | -0.1 (-2.2 to 2.0) | 1.9 (1.2 to 3.0) |
| Continuers, pregabalin vs. lamotrigine | 142 / 1,025 | -1.2 (-4.7 to 2.3) | -0.5 (-7.2 to 6.2) | 0.4 (-3.4 to 4.2) | -0.6 (-4.7 to 3.5) | 2.3 (1.0 to 5.3) |
| Mother with epilepsy, pregabalin vs. lamotrigine | 33 / 1,537 | -5.6 (-10.7 to -0.4) | -11.2 (-35.7 to 13.3) | -4.2 (-10.0 to 1.6) | 3.3 (-4.7 to 11.2) | 4.2 (1.6 to 11.4) |
| Female infants, pregabalin vs. lamotrigine | 265 / 1,146 | -2.0 (-4.6 to 0.7) | -3.2 (-8.7 to 2.2) | -1.9 (-4.2 to 0.5) | -2.4 (-5.1 to 0.3) | 1.9 (1.0 to 3.4) |
| Male infants, pregabalin vs. lamotrigine | 257 / 1,044 | -0.2 (-3.1 to 2.6) | 0.4 (-4.6 to 5.4) | -0.2 (-3.5 to 3.1) | 1.2 (-1.7 to 4.2) | 1.2 (0.6 to 2.4) |
| High vs. low dose of pregabalin | 175 / 174 | 0.6 (-2.7 to 3.9) | 0.4 (-7.0 to 7.7) | 1.1 (-1.9 to 4.2) | 1.2 (-2.3 to 4.7) | 1.1 (0.6 to 2.3) |
|  |  | Birth weight z-score |  |  |  | SGA |
| Use any time in pregnancy, pregabalin vs. lamotrigine | 528 / 2,215 | -0.1 (-0.3 to 0.0) | -0.0 (-0.3 to 0.2) | -0.2 (-0.3 to 0.0) | -0.2 (-0.4 to 0.0) | 1.3 (0.6 to 3.0) |
| Use in first trimester, pregabalin vs. lamotrigine | 489 / 2,001 | -0.2 (-0.3 to -0.0) | -0.0 (-0.3 to 0.2) | -0.2 (-0.4 to -0.0) | -0.2 (-0.5 to -0.0) | 1.3 (0.6 to 3.1) |
| Continuers, pregabalin vs. lamotrigine | 142 / 1,033 | -0.1 (-0.4 to 0.1) | -0.2 (-0.7 to 0.3) | -0.1 (-0.4 to 0.2) | -0.2 (-0.6 to 0.1) | 0.6 (0.1 to 3.0) |
| Mother with epilepsy, pregabalin vs. lamotrigine | 33 / 1,550 | 0.1 (-0.3 to 0.5) | 0.2 (-0.9 to 1.2) | 0.2 (-0.2 to 0.6) | -0.3 (-0.7 to 0.2) | Not applicable |
| Female infants, pregabalin vs. lamotrigine | 270 / 1,159 | -0.1 (-0.3 to 0.1) | 0.1 (-0.3 to 0.5) | -0.1 (-0.4 to 0.1) | -0.3 (-0.5 to 0.0) | 1.9 (0.4 to 8.3) |
| Male infants, pregabalin vs. lamotrigine | 258 / 1,056 | -0.1 (-0.3 to 0.1) | 0.0 (-0.3 to 0.3) | -0.2 (-0.4 to 0.0) | -0.1 (-0.5 to 0.2) | 1.4 (0.5 to 4.2) |

|  |  | Difference (95% CI) |  |  |  | Odds Ratio<br>(95% CI) |
| --- | --- | --- | --- | --- | --- | --- |
|  | Exposed to<br>Pregabalin/Ref<br>erence, n/n | Percentile |  |  |  |  |
|  |  | Mean | 10th | 50th | 90th |  |
| High vs. low dose of pregabalin | 177 / 176 | 0.0 (-0.2 to 0.3) | -0.3 (-0.8 to 0.1) | 0.2 (-0.1 to 0.5) | 0.2 (-0.1 to 0.6) | 1.3 (0.4 to 4.6) |
| Birth length z-score |  |  |  |  |  |  |
| Use any time in pregnancy, pregabalin vs. lamotrigine | 521 / 2,186 | -0.1 (-0.2 to 0.0) | -0.0 (-0.3 to 0.2) | -0.1 (-0.3 to 0.0) | -0.1 (-0.4 to 0.1) |  |
| Use in first trimester, pregabalin vs. lamotrigine | 484 / 1,977 | -0.1 (-0.2 to 0.0) | -0.1 (-0.4 to 0.1) | -0.2 (-0.4 to 0.0) | -0.1 (-0.3 to 0.2) |  |
| Continuers, pregabalin vs. lamotrigine | 140 / 1,018 | -0.1 (-0.4 to 0.1) | -0.1 (-0.5 to 0.4) | -0.1 (-0.4 to 0.3) | -0.2 (-0.7 to 0.2) |  |
| Mother with epilepsy, pregabalin vs. lamotrigine | 32 / 1,530 | -0.1 (-0.5 to 0.3) | -0.1 (-0.9 to 0.8) | 0.1 (-0.3 to 0.6) | -0.1 (-0.7 to 0.5) |  |
| Female infants, pregabalin vs. lamotrigine | 266 / 1,144 | -0.1 (-0.2 to 0.1) | -0.1 (-0.4 to 0.2) | -0.0 (-0.2 to 0.2) | -0.0 (-0.3 to 0.3) |  |
| Male infants, pregabalin vs. lamotrigine | 255 / 1,042 | -0.1 (-0.3 to 0.1) | 0.1 (-0.2 to 0.3) | -0.2 (-0.5 to 0.1) | -0.3 (-0.7 to 0.1) |  |
| High vs. low dose of pregabalin | 172 / 175 | 0.0 (-0.2 to 0.2) | -0.3 (-0.7 to 0.0) | -0.0 (-0.3 to 0.2) | 0.1 (-0.2 to 0.5) |  |
| Birth head circumference z-score |  |  |  |  |  | Microcephaly |
| Use any time in pregnancy, pregabalin vs. lamotrigine | 516 / 2,160 | -0.0 (-0.1 to 0.1) | 0.1 (-0.1 to 0.3) | -0.0 (-0.2 to 0.1) | -0.1 (-0.3 to 0.1) | 1.2 (0.5 to 2.9) |
| Use in first trimester, pregabalin vs. lamotrigine | 480 / 1,951 | -0.0 (-0.2 to 0.1) | 0.1 (-0.1 to 0.4) | -0.1 (-0.2 to 0.1) | 0.0 (-0.2 to 0.3) | 1.3 (0.5 to 3.4) |
| Continuers, pregabalin vs. lamotrigine | 136 / 1,012 | -0.1 (-0.3 to 0.2) | -0.1 (-0.7 to 0.6) | -0.0 (-0.3 to 0.3) | -0.1 (-0.5 to 0.2) | 5.3 (0.9 to 30.8) |
| Mother with epilepsy, pregabalin vs. lamotrigine | 32 / 1,508 | -0.0 (-0.4 to 0.4) | 0.0 (-1.0 to 1.1) | 0.1 (-0.3 to 0.4) | 0.4 (-0.5 to 1.3) | Not applicable |
| Female infants, pregabalin vs. lamotrigine | 264 / 1,128 | 0.0 (-0.2 to 0.2) | 0.3 (-0.1 to 0.6) | -0.1 (-0.3 to 0.1) | 0.0 (-0.3 to 0.3) | 1.2 (0.3 to 4.5) |
| Male infants, pregabalin vs. lamotrigine | 252 / 1,032 | -0.0 (-0.3 to 0.2) | -0.3 (-0.6 to 0.1) | 0.0 (-0.2 to 0.2) | -0.1 (-0.4 to 0.2) | 1.6 (0.5 to 5.7) |
| High vs. low dose of pregabalin | 170 / 174 | 0.0 (-0.2 to 0.2) | 0.1 (-0.2 to 0.5) | 0.1 (-0.2 to 0.4) | -0.3 (-0.8 to 0.1) | Not applicable |

AED use was ascertained at any time in pregnancy, except where noted (indented rows). Analyses on continuers are based on data from deliveries in 2006-2013. In analyses of pregabalin vs. lamotrigine, the reference was lamotrigine in the same exposure window. In dose-response analyses, the reference was the bottom tertile of mean daily dose of pregabalin (2006-2013). All results are adjusted for birth year, maternal age at delivery, education, country of origin, marital status, body mass index, smoking in current pregnancy, alcohol dependence, diabetes, hypertension, epilepsy, depression, bipolar disorder, migraine, chronic pain, and other psychiatric disorders. When the smallest cell count was < 5, we did not produce adjusted results ("not applicable").

### Levetiracetam

First appearing in this cohort in 2002, levetiracetam use increased to be the fourth most commonly used AED in 2013 (10% of infants, Figure 1). Mothers of 99% of levetiracetam-exposed infants had a diagnosis of epilepsy; 59% of infants were exposed AED polytherapy (Table 1). Common polytherapies involved lamotrigine (91 of 180 infants), carbamazepine (48), and valproic acid (33).

On average, pregnancy duration was half a day shorter (−0.5 [−2.6 to 1.6]), birth weight was 0.1 SDs lighter (−0.1 [−0.3 to 0.0] SD), length was similar (0.0 [−0.1 to 0.1] SDs), and head circumference was 0.1 SD smaller (−0.1 [−0.3 to 0.1] SD) in pregnancies and infants exposed to levetiracetam than in those exposed to lamotrigine (Table 4 and Supporting information file 3, Table S5). In women with chronic pain, levetiracetam-exposed pregnancies were longer than lamotrigine-exposed pregnancies. Most ORs for preterm delivery were slightly above 1; adjusted ORs for SGA and microcephaly were often not estimable due to cell counts below five. Infants exposed to polytherapy had reduced head circumference (−0.6 [−0.9 to −0.3] SDs on average). Exposure to an SSRI operated as an effect-measure modifier for duration of gestation, with shorter pregnancies (−11.5 [−22.3 to −0.6]) days) in women exposed to both levetiracetam and SSRIs (Supporting information file 3, Table S6). No clear dose-response relations were observed.

**Table 4.** Association Between in-Utero Levetiracetam Exposure and the Endpoints Duration of Pregnancy and Size at Birth

|  | Exposed to<br>Levetiracetam<br>/Reference,<br>n/n | Difference (95% CI)<br>At Percentile |  |  |  | Odds Ratio<br>(95% CI) |
| --- | --- | --- | --- | --- | --- | --- |
|  |  | Mean | 10th | 50th | 90th |  |
| Pregnancy duration (days) |  |  |  |  |  | Preterm birth |
| Use any time in pregnancy, levetiracetam vs. lamotrigine | 213 / 2,133 | -0.5 (-2.6 to 1.6) | -1.0 (-6.3 to 4.3) | 0.6 (-1.2 to 2.4) | 1.6 (-0.2 to 3.3) | 1.3 (0.8 to 2.3) |
| Use in first trimester, levetiracetam vs. lamotrigine | 184 / 1,938 | -0.7 (-2.9 to 1.5) | -1.7 (-7.3 to 4.0) | 0.3 (-1.8 to 2.5) | 1.8 (0.1 to 3.6) | 1.6 (0.9 to 2.8) |
| Continuers, levetiracetam vs. lamotrigine | 144 / 990 | -1.1 (-3.5 to 1.4) | -1.0 (-7.7 to 5.7) | 1.0 (-1.7 to 3.6) | 0.2 (-2.5 to 2.9) | 1.3 (0.6 to 2.6) |
| Mother with chronic pain, levetiracetam vs. lamotrigine | 52 / 536 | 2.6 (-2.1 to 7.4) | 5.3 (-5.3 to 16.0) | 2.1 (-1.9 to 6.1) | 5.0 (-0.5 to 10.5) | Not applicable |
| Polytherapy, levetiracetam vs. lamotrigine | 87 / 346 | -0.1 (-4.0 to 3.8) | -0.5 (-8.4 to 7.4) | 0.3 (-3.7 to 4.2) | -1.1 (-4.4 to 2.3) | 1.0 (0.4 to 2.7) |
| High vs. low dose of levetiracetam | 89 / 89 | -0.2 (-4.6 to 4.3) | -5.7 (-18.1 to 6.8) | -0.0 (-4.4 to 4.4) | 1.1 (-3.4 to 5.6) | 0.4 (0.1 to 1.6) |
| Birth weight z-score |  |  |  |  |  | SGA |
| Use any time in pregnancy, levetiracetam vs. lamotrigine | 215 / 2,157 | -0.1 (-0.3 to 0.0) | -0.1 (-0.4 to 0.1) | 0.0 (-0.1 to 0.2) | -0.2 (-0.4 to 0.0) | 1.3 (0.5 to 3.0) |
| Use in first trimester, levetiracetam vs. lamotrigine | 186 / 1,961 | -0.1 (-0.3 to 0.0) | -0.2 (-0.5 to 0.0) | 0.0 (-0.1 to 0.2) | -0.1 (-0.3 to 0.1) | 1.8 (0.7 to 4.3) |
| Continuers, levetiracetam vs. lamotrigine | 146 / 998 | -0.1 (-0.3 to 0.1) | -0.2 (-0.6 to 0.2) | -0.0 (-0.2 to 0.1) | -0.2 (-0.5 to 0.1) | 1.7 (0.6 to 4.5) |
| Mother with chronic pain, levetiracetam vs. lamotrigine | 51 / 546 | -0.1 (-0.4 to 0.3) | 0.2 (-0.8 to 1.3) | 0.2 (-0.2 to 0.5) | -0.4 (-0.9 to 0.1) | Not applicable |
| Polytherapy, levetiracetam vs. lamotrigine | 88 / 349 | -0.5 (-0.7 to -0.2) | -0.5 (-1.0 to 0.0) | -0.5 (-0.9 to -0.1) | -0.4 (-0.8 to 0.1) | Not applicable |
| High vs. low dose of levetiracetam | 90 / 91 | 0.1 (-0.1 to 0.4) | 0.2 (-0.7 to 1.2) | 0.1 (-0.2 to 0.4) | 0.4 (-0.1 to 0.9) | Not applicable |
| Birth length z-score |  |  |  |  |  |  |

|  | Exposed to<br>Levetiracetam<br>/Reference,<br>n/n | Difference (95% CI) |  |  |  | Odds Ratio<br>(95% CI) |
| --- | --- | --- | --- | --- | --- | --- |
|  |  | At Percentile |  |  |  |  |
|  |  | Mean | 10th | 50th | 90th |  |
| Use any time in pregnancy, levetiracetam vs. lamotrigine | 213 / 2,128 | -0.0 (-0.1 to 0.1) | -0.1 (-0.3 to 0.2) | 0.1 (-0.0 to 0.3) | -0.1 (-0.4 to 0.2) |  |
| Use in first trimester, levetiracetam vs. lamotrigine | 184 / 1,937 | -0.0 (-0.2 to 0.1) | -0.0 (-0.3 to 0.3) | 0.1 (-0.0 to 0.3) | -0.2 (-0.4 to 0.1) |  |
| Continuers, levetiracetam vs. lamotrigine | 144 / 983 | 0.1 (-0.1 to 0.2) | 0.2 (-0.2 to 0.5) | 0.1 (-0.0 to 0.3) | -0.2 (-0.4 to 0.1) |  |
| Mother with chronic pain, levetiracetam vs. lamotrigine | 51 / 536 | 0.1 (-0.2 to 0.4) | 0.4 (-0.3 to 1.1) | 0.3 (0.1 to 0.5) | 0.1 (-0.5 to 0.7) |  |
| Polytherapy, levetiracetam vs. lamotrigine | 88 / 340 | -0.2 (-0.5 to 0.0) | -0.5 (-0.9 to -0.1) | -0.0 (-0.3 to 0.3) | -0.3 (-0.7 to 0.2) |  |
| High vs. low dose of levetiracetam | 89 / 90 | 0.1 (-0.1 to 0.4) | 0.1 (-0.5 to 0.7) | 0.0 (-0.3 to 0.4) | 0.3 (-0.2 to 0.8) |  |
| Birth head circumference z-score |  |  |  |  |  | Microcephaly |
| Use any time in pregnancy, levetiracetam vs. lamotrigine | 206 / 2,103 | -0.1 (-0.3 to 0.1) | -0.1 (-0.4 to 0.2) | -0.1 (-0.3 to 0.1) | -0.1 (-0.4 to 0.1) | 1.4 (0.6 to 3.5) |
| Use in first trimester, levetiracetam vs. lamotrigine | 178 / 1,912 | -0.1 (-0.3 to 0.1) | 0.0 (-0.4 to 0.4) | -0.1 (-0.3 to 0.1) | -0.1 (-0.4 to 0.1) | 1.6 (0.6 to 4.4) |
| Continuers, levetiracetam vs. lamotrigine | 140 / 978 | -0.0 (-0.2 to 0.1) | 0.1 (-0.3 to 0.5) | -0.0 (-0.3 to 0.2) | -0.0 (-0.4 to 0.4) | Not applicable |
| Mother with chronic pain, levetiracetam vs. lamotrigine | 50 / 536 | -0.0 (-0.3 to 0.3) | -0.2 (-0.9 to 0.6) | -0.1 (-0.3 to 0.2) | -0.3 (-0.9 to 0.2) | Not applicable |
| Polytherapy, levetiracetam vs. lamotrigine | 84 / 336 | -0.6 (-0.9 to -0.3) | -0.4 (-1.0 to 0.2) | -0.8 (-1.0 to -0.5) | -0.5 (-0.9 to -0.0) | 2.7 (0.7 to 9.6) |
| High vs. low dose of levetiracetam | 87 / 89 | -0.0 (-0.3 to 0.3) | 0.4 (-0.2 to 1.0) | 0.1 (-0.3 to 0.4) | -0.4 (-1.0 to 0.2) | Not applicable |

AED, antiepileptic drug; CI, confidence interval; SGA, small for gestational age.
AED use was ascertained at any time in pregnancy, except where noted (indented rows). Analyses on continuers used data from deliveries in 2006-2013. In analyses of levetiracetam vs. lamotrigine, the reference was lamotrigine in the same exposure window. In dose-response analyses, the reference was the bottom tertile of mean daily dose of levetiracetam (2006-2013). All results are adjusted for birth year, maternal age at delivery, education, country of origin, marital status, body mass index, smoking in current pregnancy, alcohol dependence, diabetes, hypertension, epilepsy, depression, bipolar disorder, migraine, chronic pain, and other psychiatric disorders. When the smallest cell count was < 5, we did not produce adjusted results ("not applicable").

### Valproic acid

Valproic acid exposure decreased from 18% of infants in 1996 to 8% in 2013 (Figure 1). Commonly, mothers of exposed infants had a diagnosis of epilepsy (83%); 23% were on polytherapy (Table 1).

On average, valproic acid–exposed pregnancies had a duration similar to lamotrigine-exposed pregnancies (0 [-1.2 to 1.2] days), and infants were born with the same weight for gestational age (0 [-0.1 to 0] SDs) (Table 5 and Supporting information file 3, Table S7). However, we observed a gradient in which effects assessed at the 10th percentile were in the direction of the left tail (i.e., shorter pregnancies, infants lighter for gestational age) and in the direction of the right when assessed at the 90th percentile (i.e., longer pregnancies, infants heavier for gestational age). This was also true for the comparison of high versus low valproic acid doses. The association with pregnancy duration was toward longer pregnancies when the fetus was female, opposite to what was observed in pregnancies with male fetuses: the difference was 5.4 days at the 10th percentile. We observed effect-measure modification for duration of pregnancy by smoking and use of SSRIs, which resulted in valproic acid use and smoking or SSRI use being associated with shorter pregnancies (−3.1 [−6.1 to −0.2] and −3.9 [−7.7 to −0.1] days, respectively; Supporting information file 3, Table S8). Birth length did not seem to be adversely affected. Valproic acid–exposed infants had a smaller head circumference relative to lamotrigine-exposed infants, and continuers were more strongly affected (OR for microcephaly: 3.9 [1.7 to 9.0]). For all endpoints except birth length, polytherapy-exposed infants were more severely affected, with a difference in duration of 10 days at the 10th percentile. Odds ratios were generally higher for valproic acid than for other study AEDs.

**Table 5.** Association Between in-Utero Valproic Acid Exposure and the Endpoints Duration of Pregnancy and Size at Birth

|  | Exposed to<br>Valproic<br>Acid/Reference,<br>n/n | Difference (95% CI) |  |  |  | Odds Ratio<br>(95% CI) |
| --- | --- | --- | --- | --- | --- | --- |
|  |  | Percentile |  |  |  |  |
|  |  | Mean | 10th | 50th | 90th |  |
|  |  | Pregnancy duration (days) |  |  |  | Preterm birth |
| Use any time in pregnancy, valproic acid vs. lamotrigine | 985 / 2,086 | -0.0 (-1.2 to 1.2) | -1.9 (-5.3 to 1.4) | 1.0 (-0.3 to 2.3) | 1.6 (0.4 to 2.8) | 1.5 (1.1 to 2.0) |
| Use in first trimester, valproic acid vs. lamotrigine | 845 / 1,902 | -0.1 (-1.3 to 1.2) | -1.3 (-4.9 to 2.2) | 0.8 (-0.4 to 2.0) | 1.4 (0.3 to 2.5) | 1.6 (1.1 to 2.2) |
| Continuers, valproic acid vs. lamotrigine | 253 / 996 | -0.0 (-2.0 to 2.0) | -3.9 (-10.6 to 2.7) | 1.8 (-0.4 to 3.9) | 2.4 (0.7 to 4.1) | 1.7 (1.1 to 2.8) |
| Polytherapy, valproic acid vs. lamotrigine | 115 / 299 | -3.4 (-6.9 to 0.2) | -10.0 (-19.5 to -0.5) | 0.1 (-3.4 to 3.5) | 2.2 (-1.1 to 5.4) | 3.0 (1.5 to 6.2) |
| Female infants, valproic acid vs. lamotrigine | 480 / 1,094 | 0.6 (-1.1 to 2.4) | 1.6 (-2.3 to 5.5) | 1.1 (-0.6 to 2.9) | 1.7 (0.2 to 3.3) | 1.1 (0.7 to 1.8) |
| Male infants, valproic acid vs. lamotrigine | 505 / 992 | -0.7 (-2.4 to 1.0) | -3.8 (-7.6 to -0.0) | -0.1 (-1.6 to 1.5) | 0.9 (-0.6 to 2.4) | 1.9 (1.2 to 2.9) |
| High vs. low dose of valproic acid | 165 / 167 | -1.0 (-4.9 to 2.9) | -2.4 (-10.1 to 5.3) | -0.5 (-3.9 to 3.0) | 1.0 (-1.4 to 3.5) | 1.4 (0.5 to 3.4) |
|  |  | Birth weight z-score |  |  |  | SGA |
| Use any time in pregnancy, valproic acid vs. lamotrigine | 992 / 2,110 | -0.0 (-0.1 to 0.0) | -0.1 (-0.3 to 0.1) | -0.1 (-0.2 to 0.1) | 0.1 (-0.1 to 0.2) | 1.9 (1.2 to 2.9) |
| Use in first trimester, valproic acid vs. lamotrigine | 852 / 1,924 | -0.1 (-0.2 to 0.0) | -0.1 (-0.3 to 0.1) | -0.0 (-0.2 to 0.1) | 0.1 (-0.1 to 0.2) | 2.4 (1.5 to 3.8) |
| Continuers, valproic acid vs. lamotrigine | 257 / 1,004 | -0.1 (-0.3 to 0.1) | -0.3 (-0.6 to 0.1) | -0.0 (-0.2 to 0.2) | 0.3 (0.0 to 0.5) | 2.5 (1.3 to 5.0) |
| Polytherapy, valproic acid vs. lamotrigine | 116 / 302 | -0.2 (-0.5 to 0.0) | -0.2 (-0.6 to 0.2) | -0.1 (-0.4 to 0.1) | -0.4 (-0.7 to -0.1) | 2.6 (0.6 to 11.0) |
| Female infants, valproic acid vs. lamotrigine | 484 / 1,106 | -0.1 (-0.2 to 0.0) | -0.0 (-0.3 to 0.2) | -0.1 (-0.2 to 0.0) | 0.0 (-0.2 to 0.2) | 2.5 (1.3 to 5.0) |
| Male infants, valproic acid vs. lamotrigine | 508 / 1,004 | 0.0 (-0.1 to 0.1) | -0.1 (-0.3 to 0.1) | 0.0 (-0.1 to 0.2) | 0.2 (-0.0 to 0.4) | 1.5 (0.8 to 2.9) |
| High vs. low dose of valproic acid | 169 / 168 | -0.1 (-0.3 to 0.2) | -0.4 (-0.9 to 0.1) | -0.1 (-0.4 to 0.2) | 0.4 (-0.0 to 0.8) | Not applicable |
|  |  | Birth length z-score |  |  |  |  |
|  |  | Percentile |  |  |  |  |
|  |  | Mean | 10th | 50th | 90th |  |
| Use any time in pregnancy, valproic acid vs. lamotrigine | 966 / 2,083 | 0.1 (0.0 to 0.2) | 0.0 (-0.1 to 0.2) | 0.1 (-0.0 to 0.2) | 0.2 (0.1 to 0.3) |  |
| Use in first trimester, valproic acid vs. lamotrigine | 828 / 1,901 | 0.1 (-0.0 to 0.2) | 0.0 (-0.1 to 0.2) | 0.1 (-0.0 to 0.2) | 0.2 (0.1 to 0.4) |  |
| Continuers, valproic acid vs. lamotrigine | 254 / 989 | 0.1 (-0.0 to 0.3) | 0.0 (-0.3 to 0.4) | 0.2 (-0.0 to 0.4) | 0.3 (0.0 to 0.5) |  |
| Polytherapy, valproic acid vs. lamotrigine | 112 / 295 | 0.0 (-0.2 to 0.3) | 0.2 (-0.2 to 0.5) | 0.2 (0.0 to 0.4) | -0.1 (-0.5 to 0.3) |  |
| Female infants, valproic acid vs. lamotrigine | 472 / 1,091 | 0.0 (-0.1 to 0.2) | 0.1 (-0.1 to 0.3) | 0.0 (-0.1 to 0.2) | -0.0 (-0.2 to 0.2) |  |
| Male infants, valproic acid vs. lamotrigine | 494 / 992 | 0.1 (0.0 to 0.3) | 0.0 (-0.2 to 0.2) | 0.2 (0.1 to 0.4) | 0.2 (0.0 to 0.4) |  |
| High vs. low dose of valproic acid | 167 / 167 | 0.2 (-0.1 to 0.4) | 0.3 (-0.2 to 0.8) | 0.1 (-0.1 to 0.4) | 0.3 (-0.2 to 0.8) |  |
| Birth head circumference z-score |  |  |  |  |  | Microcephaly |
| Use any time in pregnancy, valproic acid vs. lamotrigine | 931 / 2,059 | -0.2 (-0.2 to -0.1) | -0.1 (-0.3 to 0.0) | -0.1 (-0.2 to -0.1) | -0.2 (-0.3 to -0.0) | 1.7 (1.0 to 2.8) |
| Use in first trimester, valproic acid vs. lamotrigine | 802 / 1,877 | -0.1 (-0.2 to -0.0) | -0.1 (-0.2 to 0.1) | -0.2 (-0.3 to -0.1) | -0.2 (-0.3 to -0.0) | 1.8 (1.0 to 3.1) |
| Continuers, valproic acid vs. lamotrigine | 252 / 983 | -0.2 (-0.3 to -0.0) | -0.2 (-0.5 to 0.1) | -0.2 (-0.3 to -0.0) | -0.2 (-0.4 to 0.1) | 3.9 (1.7 to 9.0) |
| Polytherapy, valproic acid vs. lamotrigine | 107 / 292 | -0.5 (-0.7 to -0.2) | -0.5 (-0.9 to -0.1) | -0.4 (-0.6 to -0.1) | -0.4 (-0.9 to 0.0) | 3.1 (1.0 to 9.8) |
| Female infants, valproic acid vs. lamotrigine | 458 / 1,078 | -0.2 (-0.3 to -0.0) | -0.1 (-0.3 to 0.2) | -0.2 (-0.3 to -0.0) | -0.2 (-0.3 to 0.0) | 1.8 (0.8 to 3.7) |
| Male infants, valproic acid vs. lamotrigine | 473 / 981 | -0.1 (-0.3 to -0.0) | -0.1 (-0.3 to 0.1) | -0.1 (-0.2 to 0.0) | -0.2 (-0.4 to 0.0) | 1.8 (0.9 to 3.6) |
| High vs. low dose of valproic acid | 166 / 162 | -0.2 (-0.5 to 0.0) | -0.4 (-1.0 to 0.2) | -0.1 (-0.4 to 0.2) | 0.1 (-0.3 to 0.6) | Not applicable |

AED, antiepileptic drug; CI, confidence interval; SGA, small for gestational age.
AED use was ascertained at any time in pregnancy, except where noted (indented rows). Analyses on continuers used data from deliveries in 2006-2013. In analyses of valproic acid vs. lamotrigine, the reference was lamotrigine in the same exposure window. In dose-response analyses, the reference was the bottom tertile of mean daily dose of valproic acid (2006-2013). All results were adjusted for birth year, maternal age at delivery, education, country of origin, marital status, body mass index, smoking in current pregnancy, alcohol dependence, diabetes, hypertension, epilepsy, depression, bipolar disorder, migraine, chronic pain, and other psychiatric disorders. When the smallest cell count was < 5, we did not produce adjusted results ("not applicable"). Models restricted to polytherapy compared infants exposed to valproic acid and another AED (except lamotrigine) with those exposed to lamotrigine and another AED (except valproic acid).

### Lamotrigine

Lamotrigine use in pregnancy increased over the study period from 6% in 1996 to 47% in 2013 (Figure 1). Mothers of exposed infants often had a diagnosis of epilepsy (69%); 20% of women were on polytherapy.

In dose-response analyses, pregnancies exposed to high doses were, on average, 1.8 days shorter (−1.8 [−3.8 to 0.2]) than those exposed to low doses; the OR for preterm birth was 1.3 (0.7 to 2.2) (Table 6 and Supporting information file 3, Table S7). We did not observe an association between higher doses and smaller z-scores.

**Table 6.**
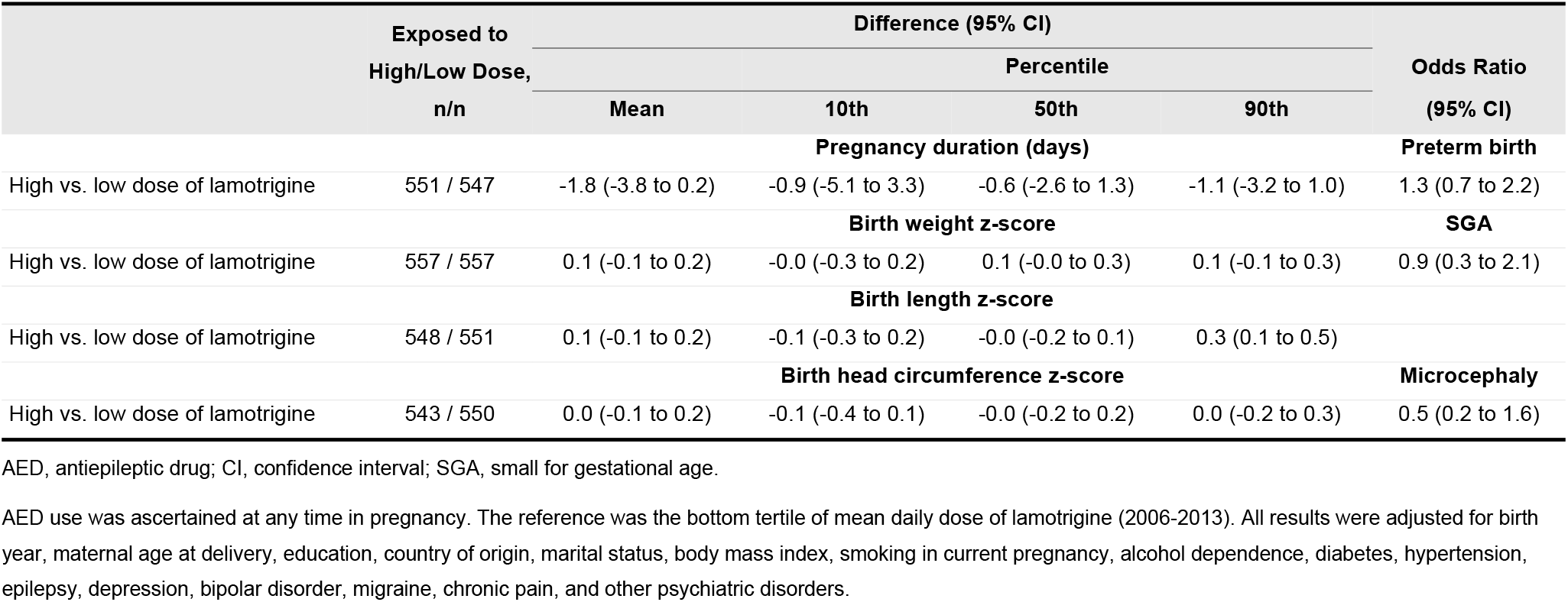
Association Between in-Utero Lamotrigine Exposure and the Endpoints Duration of Pregnancy and Size at Birth

|  | Exposed to<br>High/Low Dose,<br>n/n | Difference (95% CI) |  |  |  | Odds Ratio<br>(95% CI) |
| --- | --- | --- | --- | --- | --- | --- |
|  |  | Percentile |  |  |  |  |
|  |  | Mean | 10th | 50th | 90th |  |
| Pregnancy duration (days) |  |  |  |  |  |  |
| High vs. low dose of lamotrigine | 551 / 547 | -1.8 (-3.8 to 0.2) | -0.9 (-5.1 to 3.3) | -0.6 (-2.6 to 1.3) | -1.1 (-3.2 to 1.0) | 1.3 (0.7 to 2.2) |
| Birth weight z-score |  |  |  |  |  |  |
| High vs. low dose of lamotrigine | 557 / 557 | 0.1 (-0.1 to 0.2) | -0.0 (-0.3 to 0.2) | 0.1 (-0.0 to 0.3) | 0.1 (-0.1 to 0.3) | 0.9 (0.3 to 2.1) |
| Birth length z-score |  |  |  |  |  |  |
| High vs. low dose of lamotrigine | 548 / 551 | 0.1 (-0.1 to 0.2) | -0.1 (-0.3 to 0.2) | -0.0 (-0.2 to 0.1) | 0.3 (0.1 to 0.5) |  |
| Birth head circumference z-score |  |  |  |  |  |  |
| High vs. low dose of lamotrigine | 543 / 550 | 0.0 (-0.1 to 0.2) | -0.1 (-0.4 to 0.1) | -0.0 (-0.2 to 0.2) | 0.0 (-0.2 to 0.3) | 0.5 (0.2 to 1.6) |

AED, antiepileptic drug; CI, confidence interval; SGA, small for gestational age.
AED use was ascertained at any time in pregnancy. The reference was the bottom tertile of mean daily dose of lamotrigine (2006-2013). All results were adjusted for birth year, maternal age at delivery, education, country of origin, marital status, body mass index, smoking in current pregnancy, alcohol dependence, diabetes, hypertension, epilepsy, depression, bipolar disorder, migraine, chronic pain, and other psychiatric disorders.

### Other key variables: smoking, diabetes, and epilepsy

To put results on individual AEDs in perspective, we considered the size of the point estimates for other variables obtained from the main analysis. In all linear regression analyses for exposure at any time in pregnancy, the estimated effect of smoking was more negative than the estimated effect for all study AEDs on all study outcomes (Supporting information file 3, Table S10). For example, birth weight z-score point estimates for study AEDs were between 0 and −0.1 SDs, while, for smoking, they were between −0.4 and −0.5 SDs. Diabetes was associated with a shorter duration of pregnancy of over 1 week in analyses of all study AEDs, an effect several times larger than that of study AEDs. Point estimates for epilepsy were small or null.

## DISCUSSION

In this population-based, comparative safety cohort study involving 6,720 infants exposed to AEDs in pregnancy in Sweden during 1996-2013, we observed an increase in AED use in pregnancy over time and an evolution in preference from older to newer AEDs. With the possible exception of pregabalin, maternal characteristics were comparable across users of individual AEDs, except for the indications or uses for each drug: in the extremes, levetiracetam was used almost exclusively in women with an epilepsy diagnosis, and pregabalin was used mostly in women with chronic pain or psychiatric diagnoses. Analyses comparing individual AEDs to lamotrigine showed generally small associations (e.g., mean changes in duration of pregnancy smaller than 3 days, changes in z-scores mostly up to 0.2 SDs), which were generally milder than those observed for smoking or diabetes. Below, we contextualize our findings within what was previously known about the associations between the study AEDs and size at birth, congenital malformations and cognitive outcomes.

### Carbamazepine

On the basis of mean results from the main analysis for AED exposure at any time in pregnancy, carbamazepine-exposed infants were born 1 day earlier, were 0.1 SDs lighter and shorter, and had a head circumference that was 0.2 SDs smaller for their gestational age than infants exposed to lamotrigine; effects were dose dependent. For carbamazepine versus lamotrigine in monotherapy, our literature search identified a relative risk for SGA of 1.3 (1.0 to 1.7) [16] and an OR of 3.1 (0.9 to 10.9) [22], compared with an OR of 1.3 (0.8 to 2.0) from our study. In a myriad of statistical comparisons identified in the literature search, relative to unexposed populations, carbamazepine has been associated with shorter pregnancies and lower birth weight, length, and/or head circumference, sometimes with wide confidence intervals [13, 14, 18–21, 23, 24, 26]. Maternal exposure to carbamazepine has been associated with major congenital malformations [4] in a dose-dependent manner [5]; the association with adverse developmental, cognitive, and behavioral outcomes is less clear [8, 10].

### Pregabalin

We observed that pregabalin-exposed infants were born, on average, 1 day earlier; were 0.1 SDs lighter and shorter; and had similar head circumference for their gestational age than infants exposed to lamotrigine; no clear dose effects were seen. Because pregabalin is a relatively new AED, the literature on its safety in pregnancy is limited. Our literature search identified one study that reported elevated risk, with wide confidence intervals, for preterm delivery and SGA based on a small number of pregnancies exposed to pregabalin compared to unexposed pregnancies [41, 42]. Its association with congenital malformations is contested [11, 27, 43], and not much is known on any potential association with adverse neurodevelopmental outcomes [8].

### Levetiracetam

In our study, levetiracetam-exposed infants were born, on average, 0.5 days earlier; were 0.1 SDs lighter, with similar length; and were 0.1 SDs smaller in head circumference for their gestational age than those exposed to lamotrigine. One study identified in our literature search reported that the relative risk for the association between levetiracetam versus lamotrigine monotherapy and SGA was 1.3 (1.0 to 1.7)[16], which compares with the OR in our study for monotherapy or polytherapy combined: 1.3 (0.5 to 3.0). Comparisons with women unexposed to AEDs were less clear: one study reported that levetiracetam exposure was associated with shorter pregnancies and lighter infants [14], one reported lighter infants but practically null effects on duration of pregnancy and head circumference [20], and one reported protective effects for SGA and microcephaly [24]. The pooled risk for congenital malformations in subjects exposed to levetiracetam has been reported as similar to that for the unexposed, although some individual studies reported increased risk [4]. Developmental outcomes appear not to be negatively affected based on a single cohort [8, 10].

### Valproic acid

In our study, valproic acid–exposed infants had, on average, the same duration of gestation and birth weight for gestational age but were 0.2 SDs smaller in head circumference for gestational age than infants exposed to lamotrigine. Null mean effects masked opposite results in the two tails of the distributions of pregnancy duration and birth weight z-scores. Outcomes were worse in infants exposed to valproate in polytherapy in pregnancy, which has also been reported for congenital malformations [44]. Our literature search identified studies reporting an association of valproic acid versus lamotrigine monotherapy and SGA (relative risk: 1.5 [1.0 to 2.2] [16] and OR: 4.1 [1.1 to 15.0] [22]) that compares with that in our study (OR: 1.9 [1.2 to 3.2]). In comparison with unexposed subjects, results have been mixed: exposure to valproic acid has been reported to have a practically null effect on mean pregnancy duration [20], conferring a null [23] or increased risk for preterm delivery [14, 20]; to decrease mean birth weight [19, 20], conferring a null [23] or increased [14, 20] risk for low birth weight but not for very low birth weight [25]; to confer a lower [14, 24] or increased risk for SGA [20]; and to reduce head circumference [13, 20]. Valproic acid is a known teratogen [45], and a dose-response relation has been reported for this association [5], with variations across types of major congenital malformations [46]. In-utero exposure to valproic acid has also been reported to be associated with hearing impairment [47] and to have a dose-response relation with adverse developmental, cognitive, and behavioral effects [8, 10, 48]. In 2014, the European Medicines Agency (EMA) conducted a review on the pregnancy safety of valproic acid, after which it imposed a number of risk minimization activities in Europe [49]. Subsequent studies in France, the first country in which valproic acid was approved to treat epilepsy [50], showed that valproic acid use continued to be high [11, 51]. This triggered a second review by EMA, which then strengthened its risk minimization measures, now including a pregnancy prevention program [52].

### Lamotrigine

We observed an association between high doses of lamotrigine and shorter pregnancies (1.8 days on average). In comparisons of women exposed to lamotrigine with those unexposed, published studies reported null or adverse effects on pregnancy duration and birth weight [14, 20, 23], protective or null effects on SGA [14, 20, 24], and null effects on head circumference [13, 20]. A recent systematic review that focused on lamotrigine concluded that there was no association between lamotrigine in monotherapy and congenital malformations, preterm delivery, or SGA [28, 29]; but a dose dependency was reported for congenital malformations.[5]. Studies assessing neurodevelopmental outcomes have reported outcomes similar to those of the general population, but also a potentially increased risk for some specific deficits [8, 10].

### Secondary and sensitivity analyses, strengths, and limitations

We treated all pregnancies as independent observations because statistical models incorporating within-woman correlation would not converge; results from a sensitivity analysis including only the first infant per woman (Supporting information file 3) generally shows, as expected, wider confidence intervals. They also show some variability in point estimates, because this sensitivity analysis excluded fewer infants exposed to pregabalin but more infants exposed to valproic acid than those exposed to lamotrigine. Twelve percent of study infants had missing data, with missingness decreasing over time; the complete case analysis (Supporting information file 3) was consistent with the main analysis. We only ascertained prescriptions dispensed during pregnancy due to the lack of information on duration of use of prescribed medications; while this could have caused under-ascertainment of prescription-based exposure, we expect we captured AED use when it extended into pregnancy, from self-report during prenatal care.

Strengths of this study include our ability to incorporate exposure from both self-reports and dispensed prescriptions. Results from analyses that defined exposure based on concordant self-reports and dispensed prescriptions are consistent with the main analysis. We were able to adjust for multiple AED indications or uses and to explore associations in the tails of study outcomes. We thus identified that a zero association at the mean (i.e., results from linear regression) can mask associations at the tails of the outcome distribution, as was seen in this study for valproic acid, and duration of pregnancy and birth weight z-score using quantile regression. Another strength of this study is our ability to define our endpoints as z-scores, which we preferred because z-scores enable assessing size independently of any effect on pregnancy duration. Because other researchers may be interested in results on birth weight, length, and head circumference without this transformation, we included those results in Supporting information file 3.

We observed different effects on pregnancies with female and male fetuses for some associations, without a clear pattern. While these may reflect true effects of AEDs, they may also reflect differential fetal survival by sex perhaps in relation to sex-specific congenital malformations [53]. Table 1 shows some variation in the percentage of female infants across AEDs. We hope future research will help clarify this aspect.

The body of evidence on the associations between in-utero exposure to AEDs and maternal, pregnancy, fetal, and infant outcomes argue against combining all AEDs into a single group for safety pregnancy research. The relative prevalence of AED use in pregnancy has evolved over time, and drugs have different safety profiles, making results on the combined AEDs not comparable from one study to another and not reflective of the risk of any specific AED.

## Conclusions

We observed that commonly used AEDs have distinct safety profiles regarding duration of pregnancy and size at birth. In comparison with lamotrigine, valproic acid and carbamazepine had a more negative association with head circumference than other study AEDs. Generally, our results were of smaller magnitude for AEDs than for smoking. Associations between valproic acid and the endpoints duration of pregnancy and birth weight for gestational age in the left tail of the distributions were toward shorter pregnancies and smaller infants, although mean effects were null.

## ACKNOWLEDGMENTS

Editorial help was provided by John Forbes, and graphic services were provided by Jason Mathes, both from RTI Health Solutions (RTI-HS). Abenah Harding, from RTI-HS, provided helpful comments on a previous version of this manuscript.

## Supporting information captions

S1_Systematic_literature_search: Systematic literature search. This is Supporting information file 1.

S2_Variable_definitions: Patient characteristics and other variables-definitions. This is Supporting information file 2.

S3_Result_tables: Result tables S1 to S10. This is Supporting information file 3.

